# The antiangiogenic peptide VIAN-c4551 inhibits lung melanoma metastasis in mice by reducing pulmonary vascular permeability

**DOI:** 10.1101/2024.12.22.629954

**Authors:** Alma Lorena Perez, Magdalena Zamora, Manuel Bahena, Regina Aramburo Williams, Elva Adán-Castro, Thomas Bertsch, Jakob Triebel, Gonzalo Martinez de la Escalera, Juan Pablo Robles, Carmen Clapp

## Abstract

**Introduction:** Cancer cells drive the increase in vascular permeability mediating tumor cell extravasation and metastatic seeding. VIAN-c4551, an antiangiogenic peptide analog of vasoinhibin, inhibits the growth and vascularization of melanoma tumors in mice. Because VIAN-c4551 is a potent inhibitor of vascular permeability, we evaluated whether its antitumor action extended to a reduction in metastasis generation.

**Methods:** Circulating levels of vascular endothelial growth factor (VEGF), lung vascular permeability, melanoma cell extravasation, and melanoma pulmonary nodules were assessed in C57BL/6J mice intravenously inoculated with murine melanoma B16-F10 cells after acute treatment with VIAN-c4551. VEGF levels, transendothelial electrical resistance, and transendothelial migration in cocultures of B16-F10 cells and endothelial cell monolayers supported the findings.

**Results:** B16-F10 cells increased circulating VEGF levels and elevated lung vascular permeability 2 hours after inoculation. VIAN-c4551 prevented enhanced vascular permeability and reduced melanoma cell extravasation after 2 hours and the number and size of macroscopic and microscopic melanoma tumors in lungs after 17 days. In vitro, VIAN-c4551 suppressed the B16-F10 cell-induced and VEGF mediated increase in endothelial cell monolayer permeability and the transendothelial migration of B16-F10 cells.

**Conclusions:** These findings support the inhibition of distant vascular permeability for the prevention of tumor metastasis and unveil the anti-vascular permeability factor VIAN-c4551 as a potential therapeutic drug able to prevent metastasis generation by lowering the extravasation of melanoma cells.

## Introduction

Metastasis is the major cause of cancer death, yet metastatic activity is not the main end point of current anti-cancer treatments, and the process remains elusive [1]. A critical step in early metastasis is the exiting of tumor cells from the circulation at distant sites. Extravasation depends on the ability of tumor cells to transmigrate through the vessel wall. Tumor cells can disrupt the vascular barrier at metastatic sites by secreting vascular permeability factors [2], including VEGF, the most powerful vascular permeability stimulator [3]. The development of drugs targeting the extravasation of tumor cells offers a hopeful approach to prevent or delay the metastatic outgrowth of cancer.

VIAN-c4551 is a promising anti-cancer drug. It is a highly potent antiangiogenic cyclic heptapeptide analog of vasoinhibin [4], an endogenous antiangiogenic and vascular permeability inhibitor with significant therapeutic potential [5]. Vasoinhibin inhibits endothelial cell proliferation, migration, survival, and permeability in response to VEGF and other angiogenic and vascular permeability stimulators (basic fibroblast growth factor, bradykinin, interleukin 1β) [4,5]. Vasoinhibin helps restrict physiological angiogenesis in the retina and cartilage [6,7] and inhibits pathological angiogenesis and vasopermeability in experimental models of vasoproliferative retinopathies [8], inflammatory arthritis [9], peripartum cardiomyopathy [10], preeclampsia [11] and cancer [12]. Vasoinhibin gene transfer in prostate [13], colon [14], mammary gland [15] and melanoma [16,17] cancer cells reduce primary tumor growth and neovascularization. Furthermore, vasoinhibin gene therapy delivered two days after the intravenous inoculation of melanoma cells reduces the number and size of lung melanoma nodules [16] indicating its ability to inhibit the growth of metastasis after the engraftment of tumor cells in the metastatic region but there is no information of an action before metastatic spreading.

However, the direct use of vasoinhibin as an anti-cancer therapeutic agent is hampered by difficulties in its recombinant production [18]. These difficulties have been overcome by the development of VIAN-c4551, a vasoinhibin analog with improved pharmacological properties that conserves the efficacy and potency of vasoinhibin [4]. VIAN-c4551 inhibits VEGF-induced proliferation and permeability of endothelial cells with a potency like vasoinhibin (IC_50_=150 pM) and is orally active to inhibit melanoma tumor growth and vascularization in mice [4,19]. Because extravasation is a pivotal step of metastasis involving tumor vasopermeability factors such as VEGF [2,3] and VIAN-c4551 is a potent inhibitor of VEGF-induced vasopermeability [4,19], we aimed to investigate whether the anti-tumor action of VIAN-c4551 extended to the inhibition of the VEGF-mediated vascular permeability that mediates the extravasation and metastatic spread of melanoma cells.

## Methods

### Reagents

VIAN-c4551 (>95% pure) was synthesized by GenScript (Piscataway, NJ, USA).

### Cell culture

The mouse melanoma B16-F10 cell line (CRL-6475, ATCC, Manassas, VA, USA) expressing (B16-F10-GFP) or not the green fluorescent protein and the bovine umbilical vein endothelial cell line (BUVEC-E6E7), generated and characterized as reported [20], were cultured in high glucose Dulbecco’s Modified Eagle Medium (DMEM) (12100-038, GIBCO, Thermo Fisher Scientific, Waltham, MA, USA) supplemented with 10% fetal bovine serum (FBS) (26140-079, GIBCO) and 100 U/ml penicillin-streptomycin (L0022, Biowest, Bradenton, FL, USA).

### Vascular permeability in vitro

BUVEC-E6E7 monolayers were grown on the top of a transwell insert with 0.4 μm pores and transendothelial electrical resistance (TEER) was measured using an EVOM2 Volt/Ohm resistance meter (World Precision Instruments, Sarasota, FL, USA) as previously reported [4]. Monolayers were treated with vehicle (DMEM 10%SFB), 100 nM VIAN-c4551 or 2 µg anti-VEGF (Ranibizumab, Lucentis®, Novartis, Basel, Switzerland) for 1 hour before adding B16-F10 cells (3.5 x 10^4^ cells in 100 µl per well) or 100 µl/well of medium conditioned by B16-F10 cells (B16-F10-CM) to the luminal side. B16-F10-CM was from cells grown to 80% confluency for 48 hours. TEER was measured over a 6-hour period. VEGF levels in the conditioned medium of 80% confluent B16-F10 cells treated or not with 100 nM VIAN-c4551 were measured by the enzyme-linked immunosorbent assay (ELISA, Quantikine mouse VEGF kit #MMV00, R&D System, Minneapolis, MN, USA).

### Actin distribution

A previously described method was used [21]. BUVEC-E6E7 were seeded on coverslips placed in 24-well plates and grown to confluence. When the monolayer was completely formed, cells were treated for 1 hour with 100 nM VIAN-c4551, 2 µg anti-VEGF (Ranibizumab, Lucentis®, Novartis) or PBS followed by the addition of 100 µl B16-F10-CM or vehicle (PBS) for 1 hour. Cells were then washed, fixed with 4% paraformaldehyde for 20 minutes, permeabilized with 0.1% Triton X-100 for 15 minutes, stained for actin with 160 nM rhodamine–phalloidin (R415, Thermo Fisher Scientific) for 1 hour in darkness, and counterstained with 5 μg/ml Hoechst 33342 (B2261, Sigma-Aldrich, Saint Louis, MO, USA). Finally, the coverslips were washed, mounted, and observed under a fluorescent microscope (Olympus BX60, Tokyo, Japan).

### Transendothelial migration assay

BUVEC-E6E7 monolayers were grown on the top of a transwell insert with 8 μm pores coated with 0.38 mg/ml matrigel (354234, Corning, Corning, NY, USA). Monolayers were treated for 1 hour with 100 nM VIAN-c4551 followed by the addition of fluorescent B16-F10-GFP cells (3.5 x 10^4^ cells per well) for transendothelial migration. Conditioned medium of B16-F10 cells containing 10% FBS added to the lower (abluminal) chamber served as chemoattractant. After 16 hours, cells in the luminal side were washed and migrating cells in the abluminal side were fixed and images observed through an inverted fluorescent microscope (Olympus IX51) and quantified using the Image Pro Plus software.

### Animals

Adult female C57BL/6J mice (8-12 weeks) were housed under standard laboratory conditions (22°C; 12h/12h light/dark cycle; free access to food and water). Experiments were approved by the Bioethics Committee of the Institute of Neurobiology of the National University of Mexico (UNAM) according to the US National Research Council’s Guide for the Care and Use of Laboratory Animals (Eighth Edition, National Academy Press, Washington, D.C., USA). B16-F10 cells were trypsinized, washed and resuspended in PBS. Restrained, non-anesthetized mice were injected with 1mg/kg body weight (b.w.) VIAN-c4551 or vehicle (saline) into the lateral tail vein, 30 minutes later, 2 x 10^5^ B16-F10 or B16-F10-GFP cells resuspended in 100 µl PBS or 100 µl PBS alone were injected into the lateral tail vein. Mice for control and treatment groups were selected at random and identified with ear tags to minimize potential confounders. Groups were divided to either collect blood, evaluate pulmonary vascular permeability or quantify melanoma cell extravasation 2 hours after B16-F10 cell intravenous (i.v.) inoculation. Other groups were euthanized 17 days after B16-F10 cell injection and their lungs harvested to evaluate macroscopic and microscopic melanoma tumors. Animals were anesthetized (60% ketamine/40% xylazine; 1µl/g b.w.) before perfusion and euthanized by an overdose of ketamine/xylazine and decapitation, and all efforts were made to minimize suffering. A total of 114 animals were used, no animals were excluded. Investigators evaluating outcomes were blind to treatment assignments. Sample size was defined based on reliable differences.

### Pulmonary vascular permeability

Vascular permeability was assessed by the extravasation of albumin stained by the Evans blue dye as previously described [22] with some modifications. Briefly, the Evans blue dye (50 µl, 45 mg/kg; E2129, Sigma-Aldrich) was i.v. injected after B16-F10 cell inoculation 1 hour before perfusion. Five hundred µl of blood was withdrawn from the heart to measure Evans blue concentration in plasma, and mice were then perfused for 2 minutes via the right ventricle with 70 ml PBS (pH 3.5 at 37°C). Lungs were dried at 72°C for 24 hours and the Evans blue dye extracted by incubating each lung in 300 μl formamide (F7503, Sigma-Aldrich) for 18 hours at 72°C. Absorbance was measured in the supernatant at 620 nm using the Varioskan Flash spectrophotometer (Thermo Fisher Scientific). The dye concentration in the extracts was calculated using a standard curve of Evans blue in formamide and normalized to the lung and body weight and to the Evans blue concentration in plasma. VEGF levels in serum were measured by ELISA (Quantikine mouse VEGF kit #MMV00, R&D System).

### Lung metastasis melanoma model

Lungs were harvested and placed in Fekete’s solution as reported [23]. Tumor nodules were evaluated macroscopically on the lung surface with a stereoscope (Leica Zoom 2000, Leica Biosystems, Nussloch, Germany) and their number and size quantified using the ImageProPlus analysis software (Version 7, Media Cybernetics Inc., Rockville, MD, USA) by 2 independent operators blind to treatment. Samples were fixed in 10% formalin for 48 hours, dehydrated in ethanol, and embedded in paraffin. Seven-µm-thick lung paraffin sections from lungs were deparaffinized with xylol, rehydrated in graded alcohol series, stained with Harris’s hematoxylin and eosin solution, and digitalized using Aperio Image ScanScope (Leica) and the number and area of microscopic nodules were quantified with the ImageProPlus analysis software.

### Lung melanoma cell extravasation

Mice were perfused with 10 ml PBS for 10 minutes (1ml/minute) via the right ventricle with PBS (pH 7.4 at 37°C) to remove nonadhered melanoma cells. The left and right lungs were processed for RT-qPCR and histological evaluation, respectively. RNA was isolated using TRIzol (15596018, Invitrogen, Thermo Fisher Scientific) and retrotranscribed with the high-capacity cDNA reverse transcription kit (4368813 Applied Biosystems, Thermo Fisher Scientific). Polymerase chain reaction (PCR) products were obtained and quantified using Maxima SYBR Green qPCR Master Mix (K0223, Thermo Fisher Scientific) in a final reaction containing 20 ng of cDNA and 0.5 μM of each of the following primer pairs for murine genes: GFP fwd (5’-AAGTCGTGCTGCTTCATGTG-3’), GFP rev (5’-CAAGCTGACCCTGAAGTTCA-3’), GAPDH fwd (5’-GAAGGTCGGTGTGAACGGATT-3’) and GAPDH rev (5’-TGACTGTGCCGTTGAATTTG-3’). Amplification consisted of 40 cycles of 10 seconds at 95 °C, 30 seconds at the annealing temperature of each primer pair, and 30 seconds at 72 °C. The mRNA expression levels were calculated by the 2^−ΔΔCT^ method. The right lung was fixed in 4% paraformaldehyde for 24 hours, cryoprotected with 4% sucrose (4072-05, JT Baker, Phillipsburg, NJ, USA) for 48 hours, frozen and cryosectioned (20 µm) (Leica CM1850). Sections were observed under a fluorescent microscope (Olympus BX60) to visualize extravasated B16-F10-GFP cells.

### Statistical analysis

The overall significance threshold was set at P < 0.05. The unpaired two-tailed t-test was used for two-group comparisons and one-way or two-way ANOVA for more than two groups. After ANOVA, Tukey, Dunnett and Sidak post hoc tests were performed using GraphPad Prism version 10.4.0 for macOS (GraphPad Software, San Diego, CA, USA).

## Results

### VIAN-c4551 reduces the permeability of pulmonary vessels stimulated by B16-F10 cells

Tumor cells stimulate the vascular permeability needed for their extravasation at metastatic sites [2,24]. To evaluate melanoma cell-induced vascular leakage, mice were injected i.v. with B16-F10 cells and vascular permeability determined in lungs by the extravasation of Evans blue stained albumin at different times post-B16-F10 cell inoculation (Fig 1a). Lung vessels responded to circulating B16-F10 cells by the accumulation of the Evans blue tracer (Fig 1b) with the increase in vascular permeability being highest and lowest at 2 hours and 8 hours post-treatment, respectively (Fig 1b and c). The timing of lowest leakage (8 hours) coincided with the reported time at which most melanoma cells have extravasated following their systemic inoculation [25] and suggested 2 hours post-tumor cell injection as the window of opportunity for modifying the exit of tumor cells from the intravascular to the extravascular compartment.

**Fig 1.**
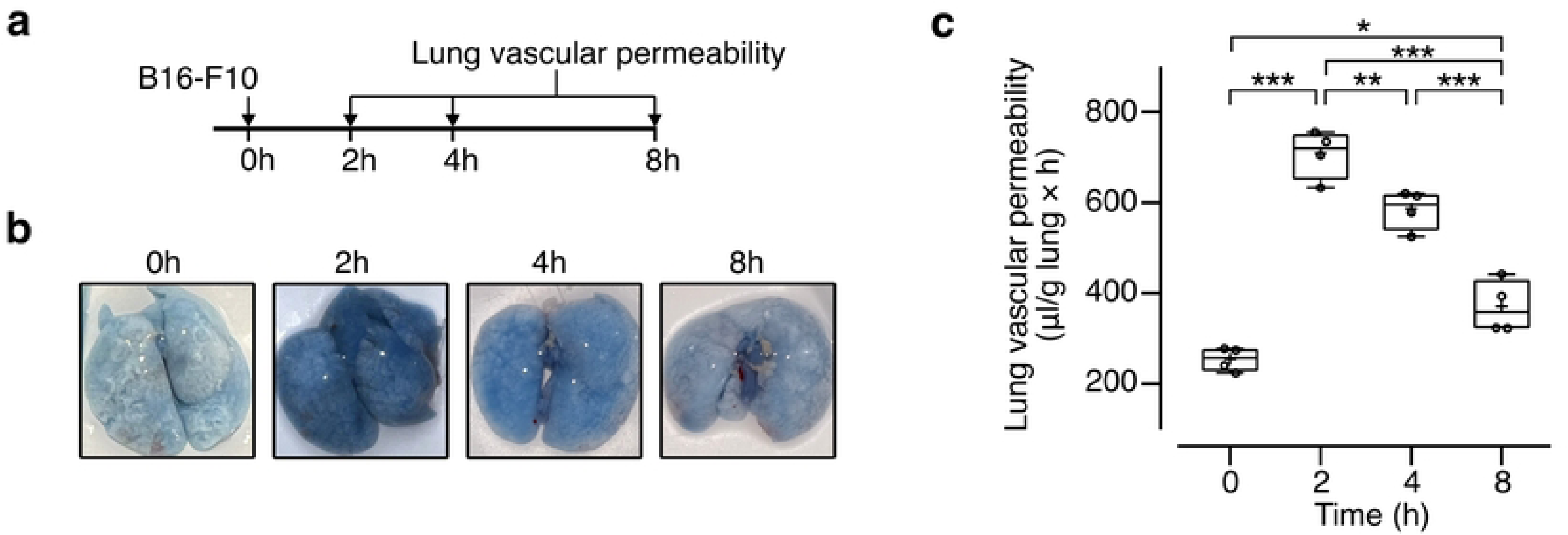
B16-F10 cells stimulate lung vascular permeability. **(a)** Timeline of the evaluation of pulmonary vascular permeability after intravenous inoculation of B16-F10 cells. **(b)** Photographs of representative lungs showing the accumulation of Evans blue-stained albumin at different times post-B16-F10 cell delivery. **(c)** Quantification of Evans blue-stained albumin as index of lung vascular permeability. Data from 2 independent experiments (n = 4) is graphed using a box and a whisker plot; the box frames the interquartile range, the horizontal line indicates the median, and the whiskers the min and max values. The mean is indicated by +. *P=0.02, **P=0.01, ***P < 0.001 (One-way ANOVA, Tukey’s multiple comparison test).

VIAN-c4551 or vehicle (saline) was i.v. injected 30 minutes before the systemic inoculation of B16-F10 cells or the i.v. injection of PBS followed by the evaluation of lung vascular leakage or serum VEGF levels after 2 hours (Fig 2a). As expected, the B16-F10 cell-induced accumulation of the Evans blue tracer was anatomically evident in lungs (Fig 2b) and accounted for a 2.5-fold increase in vascular permeability relative to PBS treated mice (Fig 2c). VIAN-c4551 prevented the upregulation of lung vascular permeability in response to melanoma cells without affecting basal vasopermeability in the absence of cells (Fig 2b and c). Because VEGF is a major vascular permeability stimulator produced by tumor cells, including B16-F10 cells [2,3], and VIAN-c4551 inhibits VEGF-induced vasopermeability [19], the circulating levels of VEGF were determined 2 hours post-tumor cell inoculation or PBS injection (Fig 2a). B16-F10 cells increased serum VEGF levels, and VEGF values were not altered by VIAN-c4551 (Fig 2d). We conclude that VIAN-c4551 abrogates lung vascular permeability not by blocking the melanoma cell release of VEGF but by inhibiting the vasopermeability action of the VEGF released by melanoma cells.

**Fig 2.**
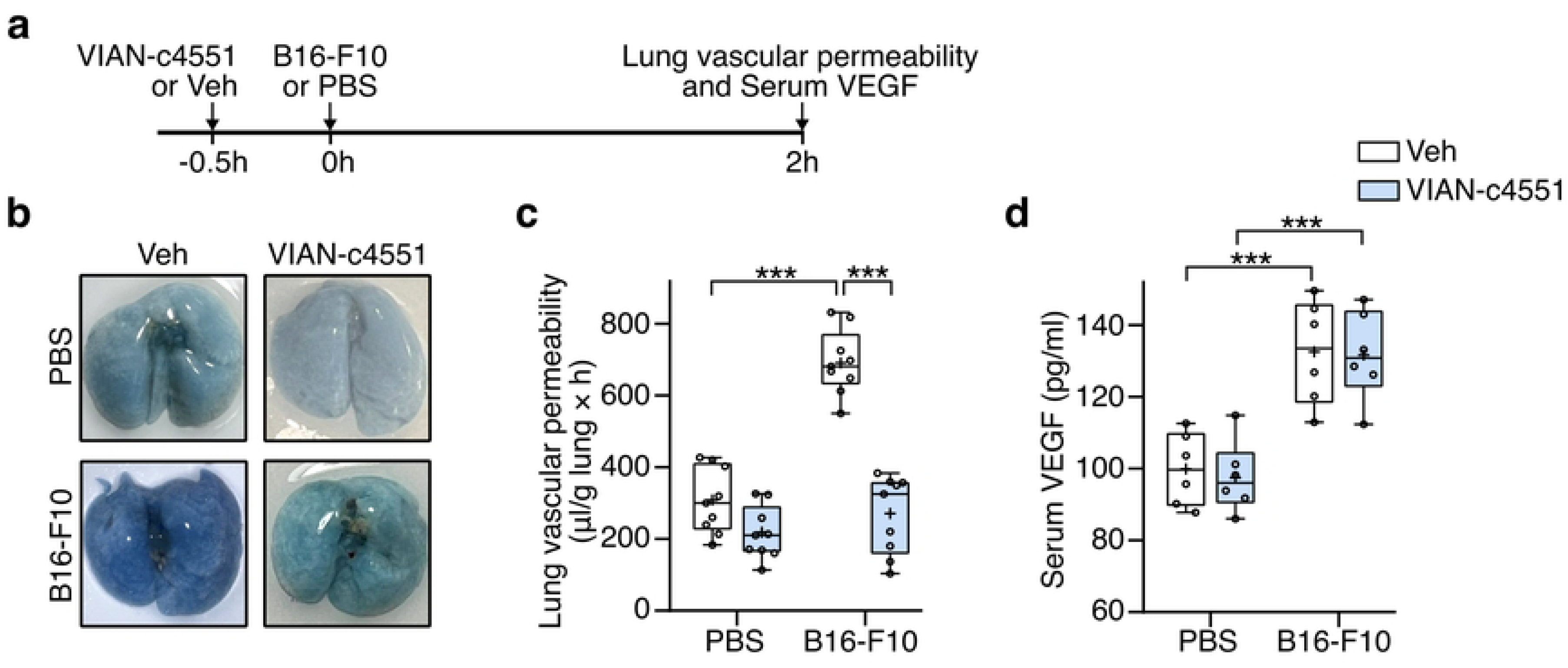
VIAN-c4551 prevents the increase of vascular permeability stimulated by melanoma cells. **(a)** Timeline of the experiment: VIAN-c4551 or vehicle (Veh) were i.v. injected 30 minutes before the i.v. delivery of B16-F10 cells or PBS. Pulmonary vascular permeability or VEGF levels were evaluated 2 hours post-tumor cells or PBS. **(b)** Photographs of representative lungs showing the accumulation of Evans blue-stained albumin as index of pulmonary vascular permeability. **(c)** Quantification of lung vascular permeability in Veh- and VIAN-c4551-treated mice after i.v. injection of PBS or B16-F10 cell inoculation. Data from 3 independent experiments (n = 9) are graphed in a box and a whisker plot; the box frames the interquartile range, the horizontal line indicates the median, and the whiskers the min and max values. The mean is indicated by +. ***P < 0.001 (Two-way ANOVA, Sidak’s multiple comparison test). **(d)** VEGF levels in serum from Veh- and VIAN-c4551-treated mice after injection of PBS or B16-F10 cell inoculation. Values are box and whiskers plot from 2 independent experiments (n =6), ***P < 0.001 (Two-way ANOVA, Sidak’s multiple comparison test).

Because a single i.v. injection of VIAN-c4551 prevented melanoma cell-induced pulmonary leakage at 2 hours, when melanoma cells are circulating [25], we hypothesized that VIAN-c4551 decreases the vascular exit of tumor cells and, thereby, reduces lung melanoma metastasis.

### A single administration of VIAN-c4551 reduces melanoma metastatic nodules in lungs

To test whether the inhibition of lung vascular leakage by VIAN-c4551 interferes with pulmonary metastasis, VIAN-c4551 or vehicle (saline) was injected i.v. 30 minutes before the systemic inoculation of B16-F10 cells and lungs were collected after 17 days to evaluate metastatic nodules (Fig 3a). Representative images showing the ventral and dorsal views of left and right pulmonary lobes showed many superficial macroscopic black nodules in vehicle-treated mice and only few in mice treated with VIAN-c4551 (Fig 3b). In agreement, VIAN-c4551 reduced by 55% the number and by 50% the size of macroscopic nodules at the lung surface (Fig 3c and d). Moreover, the histological evaluation of five longitudinal lung sections showed that VIAN-c4551 reduced by 56% and by 57% the number and size of microscopic melanoma nodules, respectively (Fig 3e-g). These findings implied that the inhibition of melanoma cell-induced vascular leakage by VIAN-c4551 is sufficient to strongly inhibit B16-F10 cell metastasis in lungs and prompted addressing this mechanism further.

**Fig 3.**
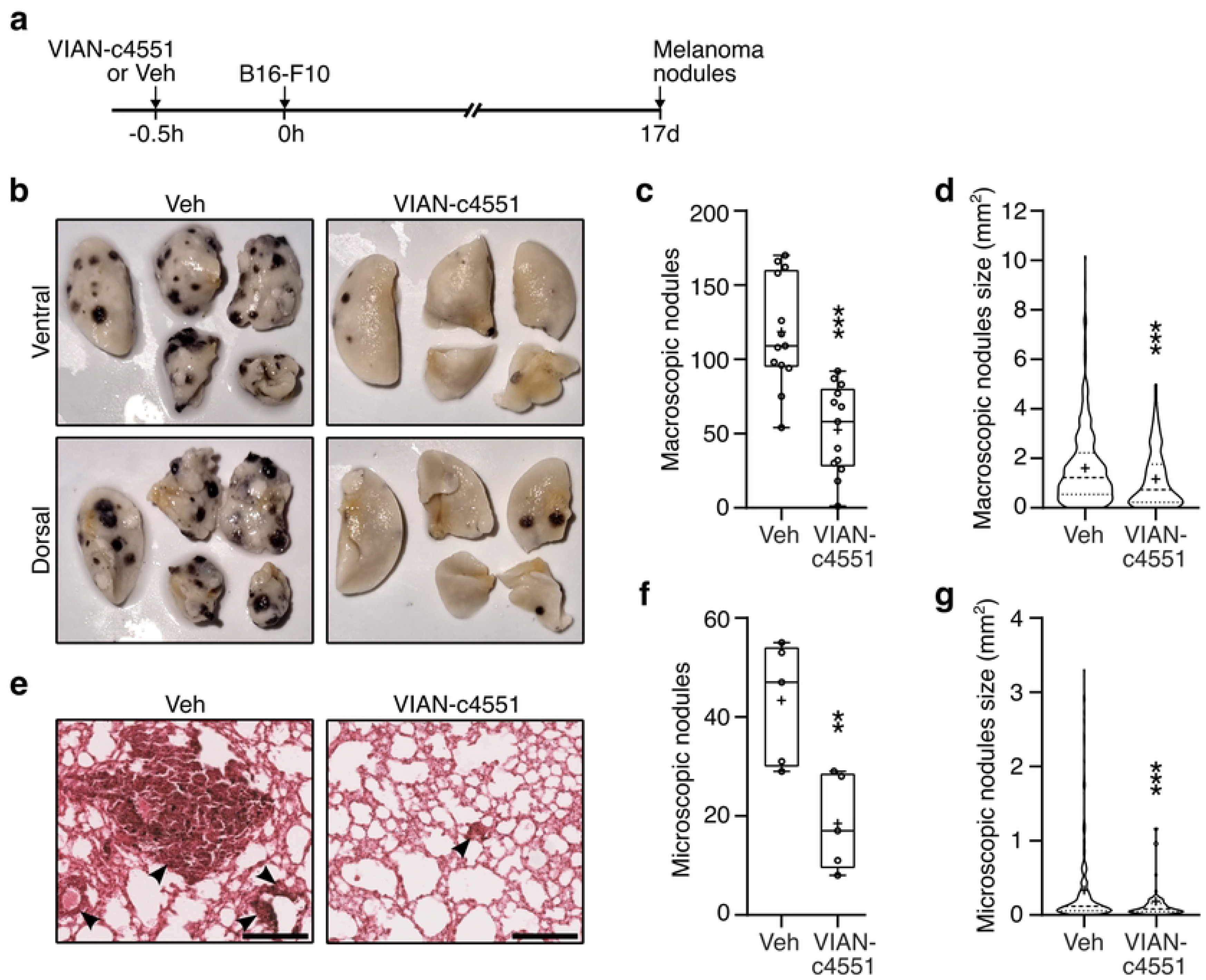
A single administration of VIAN-c4551 reduces the number and size of lung melanoma metastases in mice. **(a)** Timeline of the experiment: VIAN-c4551 or vehicle (Veh) were injected i.v. 30 minutes before the i.v. inoculation of B16-F10 cells. Lung melanoma nodules were evaluated 17 days post-tumor cell delivery. **(b)** Representative ventral and dorsal views of the left lung and the four right lobes of a mice treated with Veh or VIAN-c4551. Quantification of the number **(c)** and size **(d)** of macroscopic melanoma nodules on the lung surface. **(e)** Representative lung sections stained with hematoxylin/eosin showing microscopic melanoma nodules (arrows) (scale= 200 μm). Quantification of the number **(f)** and size **(g)** of internal microscopic melanoma nodules in lungs. Numbers of nodules are graphed in a box and a whisker plot; the box represents the interquartile range, the horizontal line indicates the median, and the whiskers the min and max values. Size is graphed in a violin plot showing median and quartiles. The mean is indicated by +. Data from 3 independent experiments (n=13). **P =0.008, ***P < 0.001 vs Veh (unpaired t-test).

### VIAN-c4551 inhibits the melanoma cell-induced permeability of endothelial cell monolayers mediated by VEGF

Transendothelial electrical resistance (TEER) of cellular monolayers measures the integrity of cell junctions and it is a quantitative indicator of the monolayer permeability [26]. B16-F10 cells (Fig 4a) and their conditioned media (B16-F10-CM) (Fig 4b) decreased the TEER of bovine umbilical vein endothelial cell (BUVEC-E6E7) monolayers throughout a 6-hour incubation period. VIAN-c4551 prevented the reduction of TEER induced by both B16-F10 and B16-F10-CM while VIAN-c4551 alone had no effect. Likewise, the anti-VEGF monoclonal fragment ranibizumab blocked the TEER reduction in response to B16-F10 and B16-F10-CM, indicating that VEGF is a substantial contributor to melanoma cell-induced hyperpermeability (Fig 4a and b). In agreement, VEGF accumulated overtime in the B16-F10-CM (Fig 4c) and, as expected from in vivo data (Fig 2d), its levels were not affected by VIAN-c4551. Increased vascular permeability due to weakened endotelial cell junctions associates with the redistribution of the actin cytoskeleton that leads to cell contraction [27,28]. The actin cytoskeleton was therefore evaluated by phalloidin-TRITC staining of BUVEC-E6E7 monolayers exposed 1 hour to the B16-F10-CM. The B16-F10-CM altered the actin cytoskeleton as revealed by increased fluorescence and frequent contracted cell bodies and these changes were prevented by VIAN-c4551 and anti-VEGF (Fig 4d).

**Fig 4.**
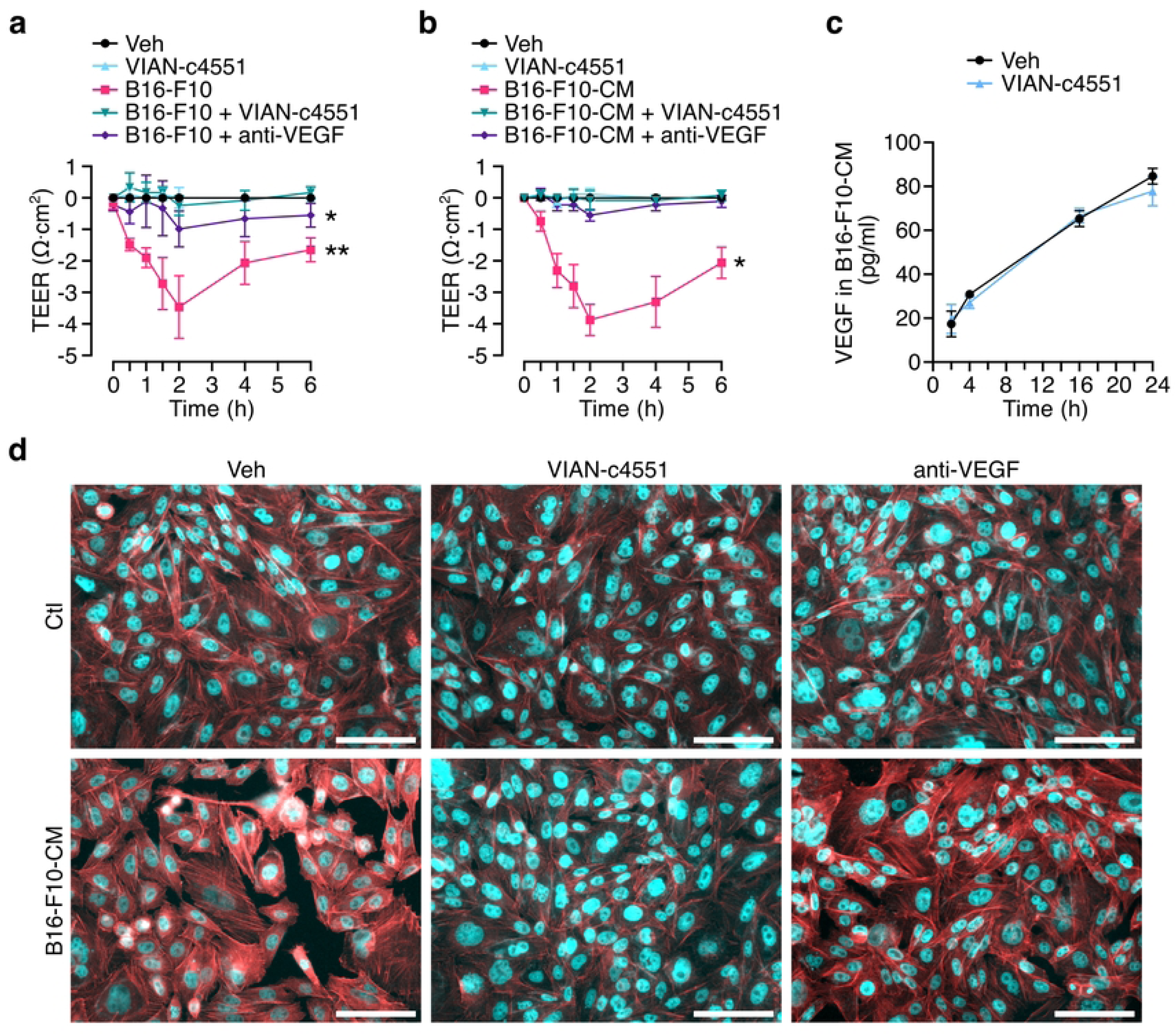
VIAN-c4551 inhibits the melanoma cell-induced permeability of endothelial cell monolayers mediated by VEGF. **(a)** Effect of B16-F10 cells on the transendothelial electrical resistance (TEER) of endothelial cell monolayers in the absence or presence of VIAN-c4551 or anti-VEGF over a 6-hour period. *P=0.0168, **P=0.0081 vs Veh. **(b)** Effect of B16-F10 conditioned media (B16-F10-CM) on TEER. *P=0.0182 vs Veh. (Repeated measurements one-way ANOVA, Dunnett’s multiple comparisons test). Values are means ± SD of 3 independent experiments. **(c)** VEGF levels in the conditioned medium of B16-F10 melanoma cells treated or not with VIAN-c4551 throughout a 24-hour incubation period. Values are means ± SD of 3 independent experiments. **(d)** Representative images of the actin cytoskeleton distribution in endothelial cell monolayers treated or not with B16-F10-CM in the absence or presence of VIAN-c4551 or anti-VEGF (scale=100 μm).

These findings supported the inhibition by VIAN-c4551 of VEGF-mediated endothelial cell permeability in response to melanoma cells and encouraged investigating whether this action interfered with the ability of B16-F10 cells to extravasate.

### VIAN-c4551 inhibits the transendothelial migration of B16-F10 cells in vitro and in vivo

The effect of VIAN-c4551 on the extravasation of melanoma cells was evaluated in vitro by measuring the migration of melanoma cells across a tight BUVEC-E6E7 monolayer. In this assay, fluorescent B16-F10-GFP cells were seeded at the top (luminal side) of a BUVEC-E6E7 confluent monolayer and the number of fluorescent cells found at the bottom (abluminal side) of the monolayer indicated transendothelial tumor cell migration. VIAN-c4551 reduced by 54% the number of B16-F10-GFP cells crossing the endothelial cell monolayer (Fig 5a and b), thereby supporting the inhibition of tumor cell extravasation by VIAN-c4551. Extravasation of melanoma cells was also studied in vivo. Mice were injected i.v. with VIAN-c4551 or vehicle (saline) 30 minutes before the i.v. inoculation of B16-F10-GFP and 2 hours later mice were perfused and lungs collected to visualize tumor cell extravasation in the premetastatic lung through the histological presence of fluorescent tumor cells (Fig 5c) or the mRNA expression levels of GFP (Fig 5d). VIAN-c4551-treated mice demonstrated lower numbers of fluorescent tumor cells and reduced lung expression of GFP relative to controls.

**Fig 5.**
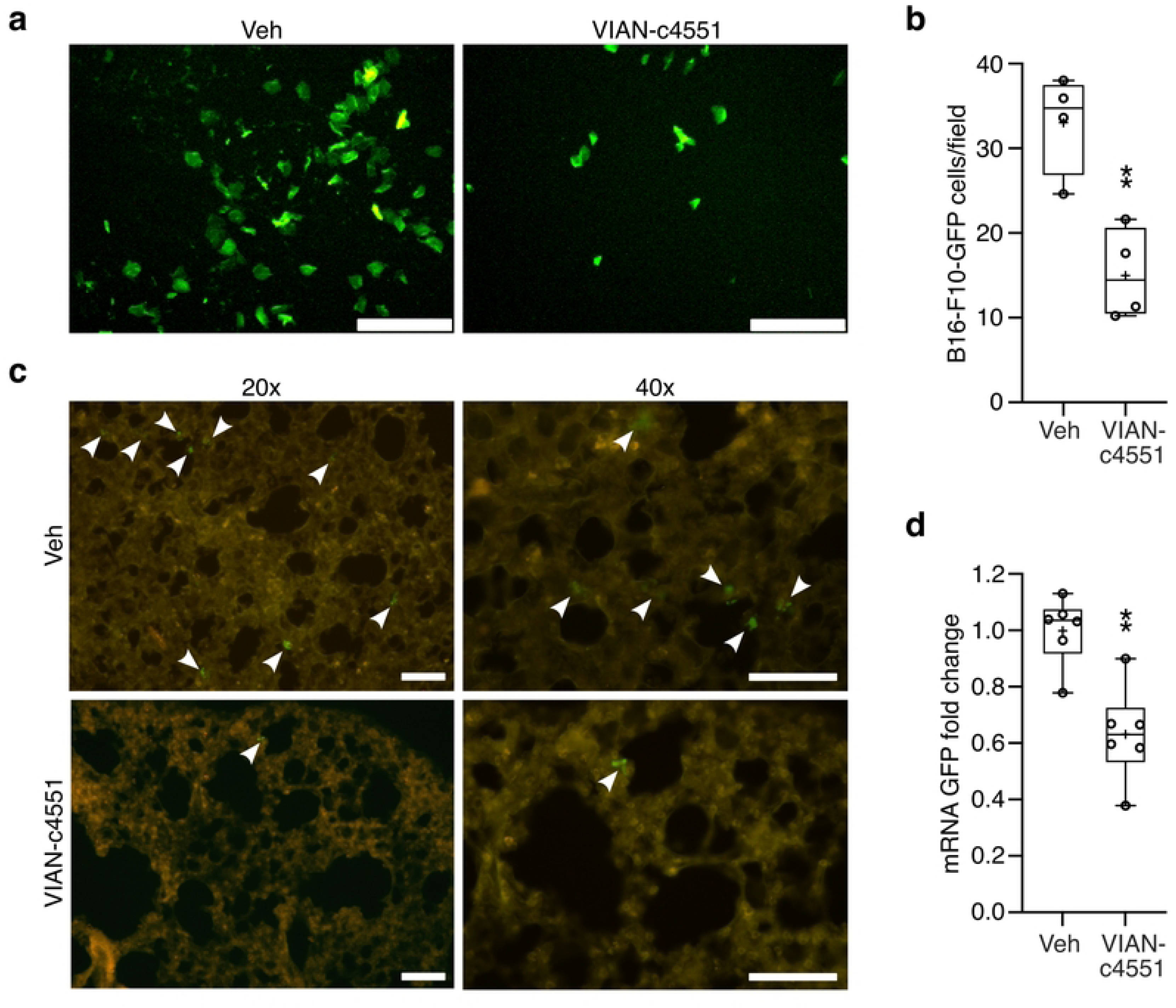
VIAN-c4551 reduces the in vitro transendothelial migration and the lung extravasation of B16-F10 melanoma cells. **(a)** Representative images showing the B16-F10 cells expressing GFP that migrated across a confluent endothelial cell monolayer in the presence of vehicle (Veh) or VIAN-c4551. **(b)** Quantification of the number of migrated B16-F10-GFP cells. Values are box and whiskers plot from 4 independent experiments. The mean is indicated by +. **P = 0.0042 vs Veh (unpaired t-test) (scale=120 μm). **(c)** Representative lung sections showing extravasated B16-F10-GFP cells 2 hours after their inoculation (arrows) (scale = 100 μm). **(d)** GFP mRNA levels in lungs from Veh- and VIAN-c4551-treated mice. Values are box and whiskers plot from 2 independent experiments (n =6). The mean is indicated by +. **P = 0.0014 vs Veh (unpaired t-test).

## Discussion

Metastasis is divided into pre-metastatic and post-metastatic phases that depend on whether tumor cells have or not engrafted at the metastatic region [2]. Because metastasis from a primary tumor can occur before primary cancer is detected [29], there is an urgent need to understand and target the premetastatic process involving the local invasion and intravasation of tumor cells at the primary site and their extravasation in remote organs [1,2]. Notably, primary tumors stimulate extravasation by increasing the permeability of blood vessels at metastatic sites [2,3,24,30,31]. Here, we support the inhibition of distant vascular permeability for the prevention of tumor metastasis and unveil the anti-vasopermeability factor VIAN-c4551 as a potential therapeutic drug able to prevent metastasis generation.

Melanoma (B16-F10) and breast (MDA-MB-231) cancer cell lines increase the permeability of lung blood vessels before metastasis by the production of vasoactive factors such as angiopoietin (Angpt) 2 and 4, matrix metalloproteinases (MMP) 1, 2, 3, and 10 and downstream effectors of VEGF (epiregulin and cyclooxygenase 2). These factors increase the permeability of lung capillaries that facilitates the transendothelial passage of tumor cells to seed pulmonary metastases [24,30,31]. Because increased vascular permeability favors extravasation, an essential step in pre-metastatic progression, inhibitors of vasopermeability represent promising therapeutics. One such inhibitor is VIAN-c4551, the cyclic heptapeptide analog of the endogenous anti-angiogenic protein vasoinhibin.

Like vasoinhibin, VIAN-c4551 inhibits the increase in vascular permeability induced by VEGF by blocking the production of nitric oxide derived from endothelial nitric oxide synthase (eNOS), an important signal that destabilizes the endothelial cell barrier through posttranslational modifications of adherens junction proteins [19,32–34]. Consistently, VIAN-c4551 and/or vasoinhibin protects against disruption of the blood retinal barrier [19,32,35] and the hypervasopermeability of the pannus [36] in experimental models of diabetic retinopathy and rheumatoid arthritis. However, beneficial effects against the extravasation of premetastatic tumors remain to be determined.

In our study, we inoculated B16-F10 cells into the tail vein of mice to circumvent primary tumor formation with the advantage of establishing a correlation between short-term lung hypervasopermeability and rapid metastatic progression. We showed that melanoma cells maximally increase pulmonary vascular permeability and extravasate at 2 hours post inoculation and that hypervasopermeability lasted at least 8 hours. The timing is consistent with the reported 8 hours required for melanoma cells to extravasate after their systemic delivery [25] and, thereby, with their metastatic seeding. Furthermore, enhanced vasopermeability at 2 hours associated with the upregulation of circulating VEGF levels suggesting the release of VEGF by melanoma cells as responsible mechanism. Accordingly, we hypothesized that the acute treatment with a VEGF inhibitor such as VIAN-c4551 would block the increase in lung vascular permeability and interfere with the extravasation process leading to fewer pulmonary metastasis. In agreement, a single intravascular injection of VIAN-c4551 inhibited the early increase in lung vasopermeability and the extravasation of melanoma cells and was enough to substantially reduce the number and size of macroscopic and microscopic metastatic nodules in lungs.

Inhibition of lung vascular leakage was previously reported to limit the number of B16-F10 cells exiting the circulation and their metastatic spread [24]. However, in contrast to our study, vasopermeability was evaluated in the presence of the primary melanoma tumor and after longer times (days) following B16-F10 cell intravascular inoculation. The authors proposed as a mechanism the altered lung microenvironment by the primary tumor through the local expression of vasopermeability factors, namely Angpt 2, MMP3, and MMP10, but not VEGF [24]. However, VEGF circulating levels were not evaluated.

The contribution of systemic VEGF to tumor cell extravasation is consistent with circulating cancer cells producing VEGF that is directly related to their metastatic potential [37] and with highly metastatic cancer cells circulating as multicellular clusters [38] whose extravasation is favored by vascular leakage. Also, the VEGF inhibitor VIAN-c4551 blocked melanoma cell-induced lung vasopermeability, melanoma cell extravasation and metastatic growth. Furthermore, our in vitro data showed that VEGF secreted by B16-F10 cells perturbs endothelial cell junctions in a manner that facilitates the passage of tumor cells across endothelial cell monolayers and that VIAN-c4551 prevented melanoma cell-induced vascular leakage like the anti-VEGF antibody fragment ranibizumab, which blocks the interaction of VEGF with its receptor [39]. Of note, VIAN-c4551 did not reduce the upregulation of VEGF circulating levels following melanoma cell inoculation nor the release of VEGF by cultured melanoma cells indicating that VIAN-c4551 targets the action and not the production of VEGF.

Our study does not rule out that other vasopermeability factors produced by melanoma cells are targeted by VIAN-c4551. Vasoinhibin is a broad acting agent that inhibits the action of different vascular permeability agents [5]. It reduces vascular leakage by blocking the phosphorylation/activation and the calcium-calmodulin binding/activation of eNOS in response to VEGF, bradykinin, acetylcholine, diabetic vitreous, and arthritic joints [32,34–36,40]. Likewise, vasoinhibin inhibits the action of several proangiogenic factors (VEGF, bFGF, bradykinin, IL-1β) with an impact on angiogenesis-dependent diseases, including cancer [12]. In fact, there is evidence that vasoinhibin inhibits the growth and vascularization of melanoma metastatic nodules in lungs during the post-metastatic phase [16], i.e., after pulmonary engraftment and colonization [1,25]. In that study [16], an adenoviral vector encoding the vasoinhibin isoform of 16 kDa (16k PRL) delivered 2 days post-melanoma cell intravascular inoculation, reduced the number and size of lung metastases. The tardy 2-day treatment and the delayed upregulation of vasoinhibin by gene therapy precluded evaluating the influence of vasoinhibin at the pre-metastatic phase.

A major obstacle for addressing vasoinhibin action in animal models has been the lack of sufficient functional protein. Post-translational modifications, protein folding, and instability complicate the recombinant production of vasoinhibin [18] and the delivery of sufficient quantities of vasoinhibin needed the gene therapy approach [16,41]. These difficulties have been overcome by the development of VIAN-c4551, a potent, easy-to-produce vasoinhibin analog [4]. VIAN-c4551 uncovered inhibition of vascular leakage as a novel mechanism by which vasoinhibin could block cancer cell extravasation at the pre-metastatic phase. However, the effect may be stronger in VIAN-c4551 because it lacks the ability of vasoinhibin to upregulate adhesion molecules in endothelial cells [42,43] that favor melanoma-endothelial cell interaction for extravasation [44].

In conclusion, our in vitro and in vivo studies support vascular leakage as a limiting step for cancer cell extravasation and disclose VIAN-c4551 as a potential anti-cancer agent for the early prevention of metastatic spread that warrants further research.

## Acknowledgments

The authors thank Fernando López Barrera, Xarubet Ruíz Herrera, Adriana González Gallardo, Ericka A. de los Ríos Arellano, Nydia Hernández Ríos, Moisés Mendoza Baltazar, Alejandra Castilla León, María A. Carbajo Mata, Eugenia Ramos Aguilar and Martín García Servín for their excellent technical assistance.

## Notes

### Competing Interest Statement

JPR, MZ, TB, JT, GME, and CC are inventors of a submitted patent application (WO/2021/098996), which is owned by the Universidad Nacional Autónoma de México (UNAM) and JT and TB. JPR is the CEO and founder of VIAN Therapeutics Inc. MZ and CC are consultants for VIAN Therapeutics. Inc.

